# Polycystin-2 (TRPP2) Regulates Primary Cilium Length in LLC-PK1 Renal Epithelial Cells

**DOI:** 10.1101/2020.02.24.962860

**Authors:** Paula L. Perez, Noelia Scarinci, Horacio F. Cantiello, María del Rocío Cantero

## Abstract

Polycystin-2 (PC2, TRPP2) is a Ca^2+^ permeable non-selective cation channel whose dysfunction generates autosomal dominant polycystic kidney disease (ADPKD). PC2 is present in different cell locations, including the primary cilium of renal epithelial cells. Little is known, however, as to whether PC2 contributes to the structure of the primary cilium. Here, we explored the effect(s) of external Ca^2+^, PC2 channel blockers, and *PKD2* gene silencing on the length of primary cilia in wild type LLC-PK1 renal epithelial cells. To identify primary cilia and measure their length, confluent cell monolayers were fixed and immuno-labeled with an anti-acetylated α-tubulin antibody. Although primary cilia length measurements did not follow a Normal distribution, data were normalized by Box-Cox transformation rendering statistical difference under all experimental conditions. Cells exposed to high external Ca^2+^ (6.2 mM) decreased a 13.5% (p < 0.001) primary cilia length as compared to controls (1.2 mM Ca^2+^). In contrast, the PC2 inhibitors amiloride (200 μM) and LiCl (10 mM), both increased primary ciliary length by 33.2% (p < 0.001), and 17.4% (p < 0.001), respectively. *PKD2* gene silencing by siRNA also elicited a statistically significant, 10.3% (p < 0.001) increase in primary cilia length, as compared to their respective scrambled RNA transfected cells. The data indicate that maneuvers that either regulate PC2 function or gene expression, modify the length of primary cilia in renal epithelial cells. Proper regulation of PC2 function in the primary cilium may be essential in the onset of mechanisms that trigger cyst formation in ADPKD.

**Significance Statement:** Polycystin-2 (PC2, TRPP2) is a Ca^2+^ permeable non-selective cation channel causing the autosomal dominant polycystic kidney disease (ADPKD). The importance of intact cilia and of fully functional polycystins in the onset of ADPKD cyst formation, point to yet unknown signaling mechanisms occurring within this organelle. We determined that the extracellular Ca^2+^ concentration, PC2 channel blockers, and *PKD2* gene silencing, all contribute to the length of primary cilia in wild type LLC-PK1 renal epithelial cells. The data indicate that proper regulation of PC2 function in the primary cilium may be essential in the onset of mechanisms that trigger cyst formation in ADPKD.

## Introduction

ADPKD is caused by mutations in either of two genes, *PKD1* or *PKD2* and is characterized by the development of epithelial-lined cysts in various organs, chiefly including the kidney and the liver.^1^ The protein products of *PKD1* and *PKD2* are transmembrane proteins called polycystin-1 (TRPP1, PC1) and polycystin-2 (TRPP2, PC2), respectively, whose localization to primary cilia was an important clue in directly linking primary cilia to renal cysts.^2^ It is now know that the vast majority of mutated proteins linked to renal cystic diseases can be localized to the primary cilium or its associated structures such as the basal body, centrosomes or ciliary transition zone.^3^

Primary cilia transduce different signals from extracellular stimuli regulating cell proliferation, differentiation, transcription, migration, polarity, and survival.^4^ Many receptors present in the primary cilium are necessary to recognize specific hormones such as somatostatin, growth factors or morphogens such as Sonic hedgehog (Shh) and Wnt, which play essential roles in the embryonic phase.^5^ Recently, a direct relationship has been found between the morphology of the primary cilium and the pathogenesis of several diseases known as ciliopathies, among which are retinitis pigmentosa, ADPKD and autosomal recessive polycystic kidney disease (ARPKD), abnormalities in axial embryonic asymmetry, obesity, cancer and the Bardet-Biedl and the Orofacial-digital type 1, syndromes.^5–7^ Further, changes in primary cilium length occur in tissues and cells grown under various physiological and pathological conditions.^8–11^ It has been shown, for example, that the length of renal primary cilia increases in ischemic mouse kidneys and in human kidney transplants that suffer acute tubular necrosis.^11, 12^ Abnormally long primary cilia have also been associated with juvenile cystic kidney disease.^10^ Conversely, in the mouse model with partial loss of the ciliary protein polaris, the animals died of ARPKD and the kidneys showed abnormally short primary cilia.^9^ ADPKD in particular is thought to develop by dysfunction of a PC1/PC2 functional receptor/ion channel complex present in various cell locations, including the primary cilium.^13, 14^

PC2 is a Ca^2+^-permeable non-selective cation channel of the TRP channel superfamily.^15^ Thus, PC2 is involved in Ca^2+^ entry steps of different epithelial tissues and organs.^15–19^ PC2 contribution to cell signaling and Ca^2+^ homeostasis has also been reported in different cellular compartments, including the primary cilium.^17, 20, 21^ The mechanosensor hypothesis of primary cilium function was introduced to suggest that changes in fluid flow outside the cell may be recognized and transduced by primary cilia to elicit cell signaling.^22, 23^ A functional PC2 channel was also reported in primary cilium.^24^ However, the mechanosensory properties of primary cilia in kidney cells have recently been questioned along with PC1’s function as a flow sensor,^23^ because no ciliary Ca^2+^ influx was observed consistent with mechanosensitive channel activation.^25^ It was thus suggested that if the mechanosensory ability of primary cilia do exist, it could induce mediators and not initiate cell Ca^2+^ signaling.^14^ The importance of both intact cilia and fully functional polycystins in the onset of ADPKD cyst formation point to unknown signaling mechanisms occurring within cilia called “cilia-dependent cyst activation”.^14^ It is presently uncertain as to whether an exclusive the so ‘cilia hypothesis’ can explain the highly variable renal phenotypic spectrum seen in different diseases.^26^

In context of the onset of the ADPKD ciliopathy in particular, little is yet known as to the possible link(s) between PC2 channel function and the morphology of the primary cilium. Therefore, in the present study, we evaluated the effect of different maneuvers to impair either the function and/or expression of PC2 on the length of the primary cilium of LLC-PK1 renal epithelial cells. The effect(s) of a high external Ca^2+^concentration, the PC2 inhibitors Li^+^ and amiloride, and the inhibition of *PKD2* gene expression were assessed on the length of the primary cilium. The results indicated that a functional PC2 is a key element in the regulation of the length of the primary cilium in LLC-PK1 renal epithelial cells, which may help explain the initial events in cyst formation and the onset of ADPKD.

## Methods

### Cell culture and immunochemistry

Wild-type LLC-PK1 renal cells (ATCC) were cultured as previously described,^24^ in Dulbecco’s modified Eagle’s medium (DMEM) supplemented with 3% fetal bovine serum (FBS), without antibiotics. Cells were seeded onto glass coverslips, and grown at 37°C, in a humidified atmosphere with 5% CO_2_ to reach full confluence in two-to-three weeks in culture. Confluent cells were used for immunocytochemical studies (Fig. 1a).

**Fig. 1:**
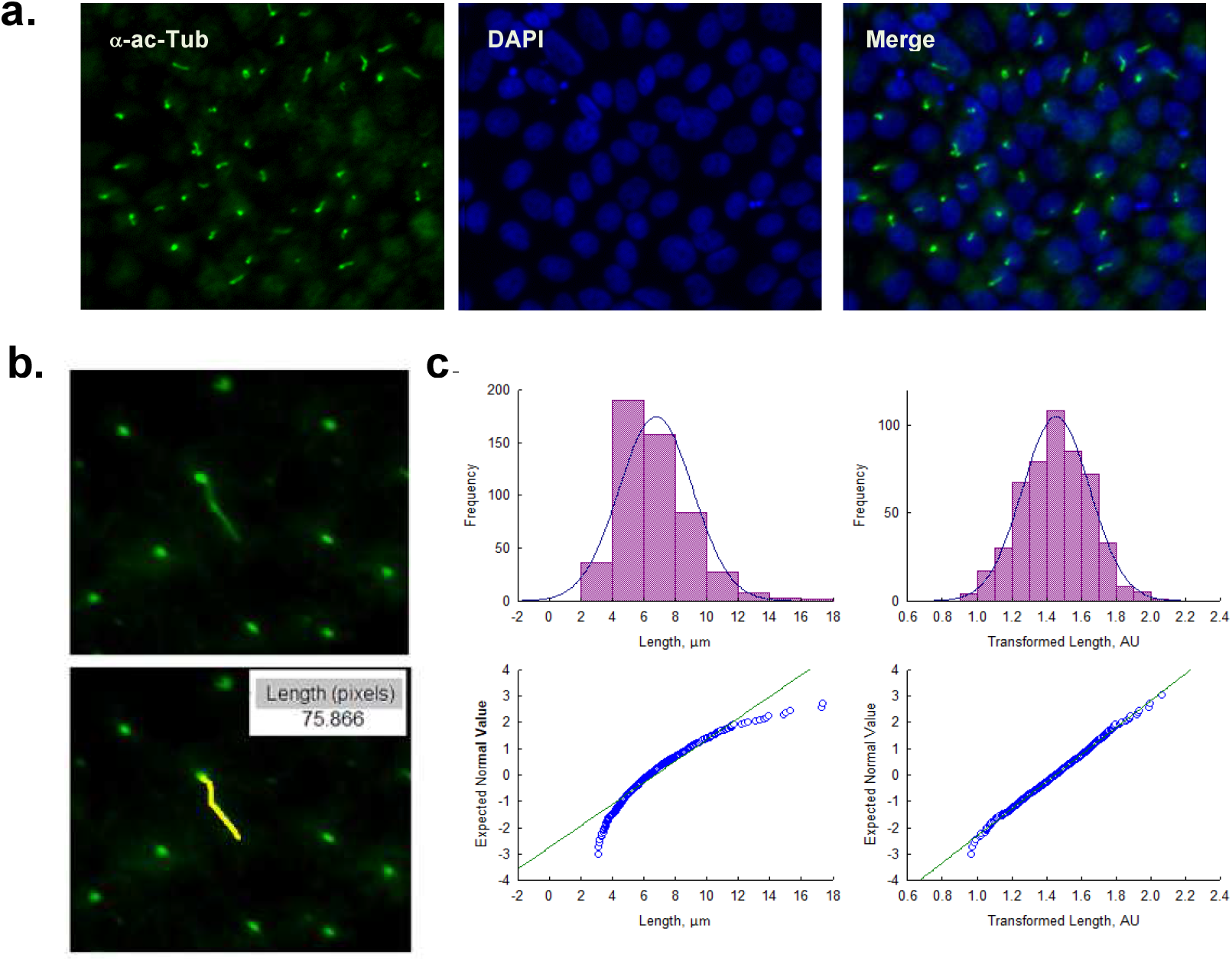
Primary cilia inmunolabeling and measurement. **a.** Confluent monolayers of wild-type LLC-PK1 cells were labeled with an anti-acetylated α-tubulin antibody (Green, Left Panel), and DAPI (Blue, Middle Panel). The merged image is shown on the Right. **b.** Primary cilia were identified (x40) and measured with the IPLab software. **c.** *Top.* Data distribution (Frequency) histograms before (Left panel) and after the Box-Cox transformation for the variable “Length of the primary cilium” are shown before and after Box-Cox transformation. The symmetry reached after the transformation is consistent with a Normalized distribution. *Bottom*. Probability density function plot before (Left) and after Box-Cox transformation. Transformed data points (Blue) are considerably closer to the fitted Normal distribution line (Green).

### Immunocytochemistry

Confluent cell monolayers exposed overnight to the various experimental conditions were rinsed twice with phosphate-buffered saline (PBS), and fixed for 10 min in a freshly prepared solution of para-formaldehyde (4%) and sucrose (2%). Cells were then washed three times with PBS, blocked for 30 min with BSA (1%) in PBS and incubated for 60 min with anti-acetylated-α-tubulin antibody (Santa Cruz Biotechnology) to identify primary cilia.^24^ Cells were counter-stained with DAPI to locate cell nuclei and mounted with Vectashield mounting medium (Vector Laboratories, Burlingame, CA). Cells were viewed under an Olympus IX71 inverted microscope connected to a digital CCD camera C4742-80-12AG (Hamamatsu Photonics KK, Bridgewater, NJ). Images were collected and analyzed with the IPLab Spectrum acquisition and analysis software (Scanalytics, Viena, VA), running on a Dell-NEC personal computer.

### Reagents

Unless otherwise stated, chemical reagents, including CaCl_2_, LiCl, amiloride, and EGTA were obtained from Sigma-Aldrich (St. Louis, MO, USA), and diluted at their final concentrations as indicated.

### Free Ca^2+^ calculations and Ca^2+^ chelation

The nominal free-Ca^2+^ solution was prepared as follows: the Ca^2+^ chelating agent EGTA (ethylene-bis(oxyethylenenitrilo) tetraacetic acid, 100 mM) was dissolved in NaOH and titrated with HCl to reach pH ~7.0, and used at a 1 mM final concentration. The final Ca^2+^ concentration was calculated by:

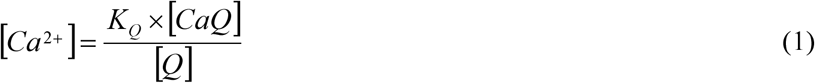

where *K*_*Q*_, is the dissociation constant of the Ca^2+^-chelator complex, [*Q*] is the concentration of the free chelating agent, and [*CaQ*] is the concentration of Ca^2+^ bound to *Q*. The nominal Ca^2+^ free solution was estimated to have a Ca^2+^ concentration of 0.32 nM. Whenever indicated, CaCl_2_ was added from a 500 mM stock solution to reach either free Ca^2+^ 1.2 mM (normal) or 6.2 mM (high) free Ca^2+^. The solutions were labeled throughout as 0 Ca, 1.2 Ca, and 6.2 Ca, respectively.

### PKD2 gene silencing

Silencing of *PKD2* gene expression in cultured LLC-PK1 cells was conducted using the small interfering RNA technique, as recently reported.^27^ Briefly, two 21-nt PKD2-specific synthetic siRNAs, one of which was a fluorescent (fluorescein) probe, were synthesized by Invitrogen (Buenos Aires, Argentina), as well as a 19-nt irrelevant sequence as scrambled control (Ir-siRNA). All constructs bore dTdT overhangs at the 3’ end, as originally reported.^28^ The siRNAs sense sequences were as follows; (siPKD2) GCUCCAGUGUGUACUACUACA, starting at 906 in exon 3 of the porcine PKD2 gene, and (Ir-siRNA) UUCUCCGAACGUGUCACGU, as scrambled control. siRNA transfection was conducted as follows:^27^ cell cultures were trypsinized and placed at 70% confluence in 35-mm cell culture dishes containing DMEM supplemented with 3% FBS at 37°C in a 5% CO_2_ atmosphere. The following day, transfection was performed with Lipofectamine 2000 (Invitrogen). Tubes were added either scrambled (Irss, 10 μl) or antisense (siPKD2, 10 μl) RNA with Optimum medium (100 μl) in the absence or presence of Lipofectamine (2 μl). Briefly, the tubes were incubated for 5 min at room temperature and then mixed with either Irss or siPKD2 for another 20 min (200 μl total volume). Incubation was conducted by medium change with a mixture of fresh medium (800 μl, DMEM plus 3% FBS), and 200 μl of the transfection mixture. The total transfection time was three overnights (72 h). Silencing efficiency was confirmed by Western blot technique, as previously reported.^27^

### Measurement of primary cilia length

The length of extended primary cilia on the fixed confluent monolayer was manually measured from 2D images with the image analysis program ImageJ (NIH software), as described elsewhere.^8, 29, 30, 31^ Briefly, FITC-labeled fluorescent primary cilia were traced with the “Freehand Line” tool of the software to obtain its length in pixels (Fig. 1b). The results were then converted into μm by calibration with a Neubauer chamber (Hausser Scientific, Horsham, PA, USA). Although this technique risks a selection bias, it is considered as reliable, allowing the measurement of a large number of irregular primary cilia, which facilitates their subsequent statistical analyses (see Supplemental Material).

### Statistical analyses

Experimental data, usually *n* > 150 measurements/experiment under each condition were collected from three to four repeats (*N* experiments). Under all tested experimental conditions primary cilia length measurements followed a non-Normal distribution (Figs. 1–4), also showing a large dispersion among experiments (e.g. range of 3.40 – 11.2 μm length for control condition with 1.2 mM Ca^2+^). Thus, data were first analyzed by non-parametric one-way ANOVA test performed by Kruskall-Wallis ranges, to assess differences for within each, and among different conditions. Individual experiments showed similar general tendencies among experimental conditions, although some did not support statistical significance. In such cases, individual experimental differences were further tested by the Mann-Whitney U test. Most often this approach helped improve statistical significance, but not for all the experiments in the same condition. To further improve statistical analysis, experimental data were finally processed by Box-Cox transformation,^32^ to aid in later post-hoc appropriate parametric statistical tests that required Normality and allowed further testing with the means and SEs of the data. The Box-Cox transformation is defined as a continuous function that varies with respect to the lambda (*λ*) power.^33^ Version 8.0 STATISTICA was used to implement these transformations and normalize the variable “length of primary cilia” (Transformed Length, AU). The program searched for the appropriate *λ* by maximum likelihood, such that the error function was minimal. The Box-Cox transformation function used in the present study was:

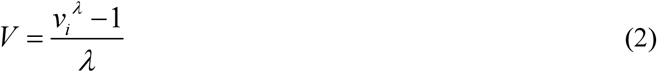

Where *V* is the result variable of the transformation, *v*_*i*_ represents the variable to be transformed and *λ* is the transformation parameter. The transformed histograms showed a remarkable change in their symmetry, approaching a Gaussian distribution (Figs. 1–4). The Normality of the data was also contrasted by comparing the empirical distribution with Normal distribution graphs (Fig. 1–4). Following data transformation, corrected mean ± SE values were obtained for each experimental condition. Transformed data were compared by one-way parametric ANOVA, and subsequent application of the Tukey post-hoc test. Statistical significance of the averaged data was accepted at p < 0.05. The Supplemental Material summarizes the statistical analyses that rendered the final comparison of mean ± SEM for the conditions control, normal Ca^2+^ (1.2 mM), and high Ca^2+^ (6.2 mM) conditions.

## Results

### Effect of external Ca^2+^ on primary cilia length

To evaluate the effect of external Ca^2+^ on the length of primary cilia, confluent monolayers of wild type LLC-PK1 cells were exposed overnight to solutions containing either 1.2 mM (normal) or 6.2 mM (high) Ca^2+^ (Fig. 2a). Cells were fixed and stained (Fig. 1a), and primary cilia length was measured as indicated in Materials & Methods (Fig. 1b). Data did not follow a Normal distribution for any of the conditions tested (Fig. 2b, Left). Values were then normalized using the Box-Cox transformation and compared among groups (Fig. 2b, Right). See the Supplemental Material for details of the statistical analysis. Cells exposed to high (6.2 mM) external Ca^2+^ had a 13.53 ± 1.25% reduction in primary cilia length as compared to the control condition in normal Ca^2+^ (4.08 ± 0.06 μm, n = 653, *vs.* 4.72 ± 0.05 μm, n = 510, p < 0.001, respectively) (Fig. 2a & b). Thus, changes in external Ca^2+^ regulated the length of primary cilia in LLC-PK1 cells. Although overnight exposure of cells to a Ca^2+^-free medium also had a decrease in primary cilia length (data not shown). The incubation procedure also produces a dramatic change in cell morphology, suggesting that the phenomenon could occur by mechanisms other than those observed with high Ca^2+^.

**Fig. 2:**
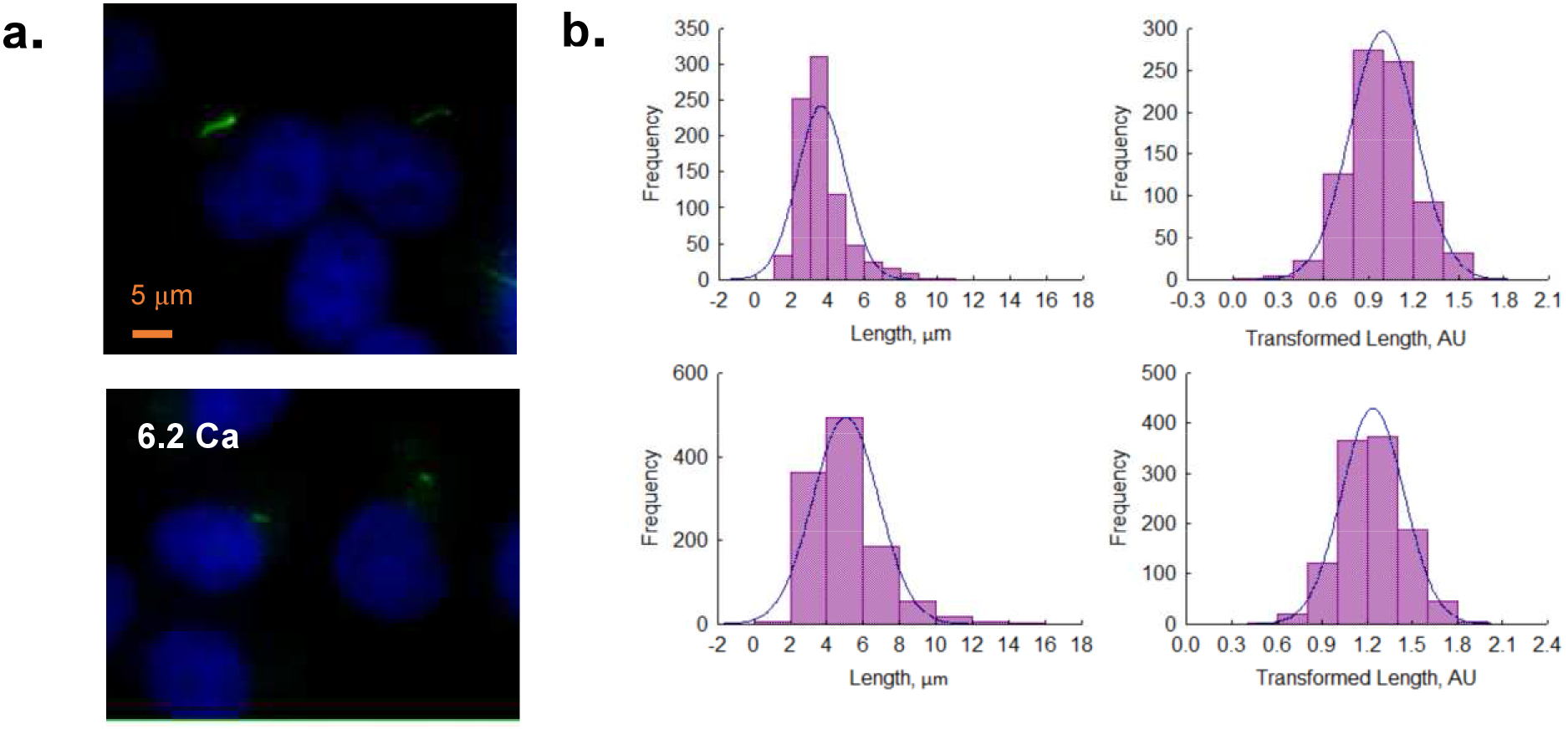
Effect of extracellular Ca^2+^ on the length of the primary cilium of wild type LLC-PK1 cells. **a.** Acetylated-α-tubulin immunolabeling of confluent of LLC-PK1 cell monolayers was used to observe the primary cilium at normal (1.2 Ca, Top panel),and high (6.2 Ca) external Ca^2+^ concentrations. Merged images also show cell nuclei by DAPI labeling (Blue). **b.** Histograms of ciliary length measurements in normal (1.2 mM, Top) and high (6.2 mM, Bottom) Ca^2+^ concentration are shown before (Left), and after (Right), Box-Cox transformation. A not Normal left-skewed distribution of data is observed before transformation. Please note that Box-Cox transformed histograms require re-transforming the values to render primary cilia length in μm (see Materials & Methods).

### Effect of amiloride on the length of primary cilia

In order to explore whether PC2 function may be implicated in the regulation of primary cilia length, the PC2 channel blocker,^15^ amiloride was tested on the LLC-PK1 cells. Confluent wild type LLC-PK1 cells were exposed overnight to a serum-free medium containing normal Ca^2+^ and 200 μM amiloride (Fig. 3a). Cells exposed to amiloride had a consistent and statistically significant 33.24 ± 1.15% increase in primary cilia length (6.29 ± 0.05 μm, n = 509 vs. 4.72 ± 0.05 μm, n = 510, p < 0.001, Fig. 3b & c).

**Fig. 3:**
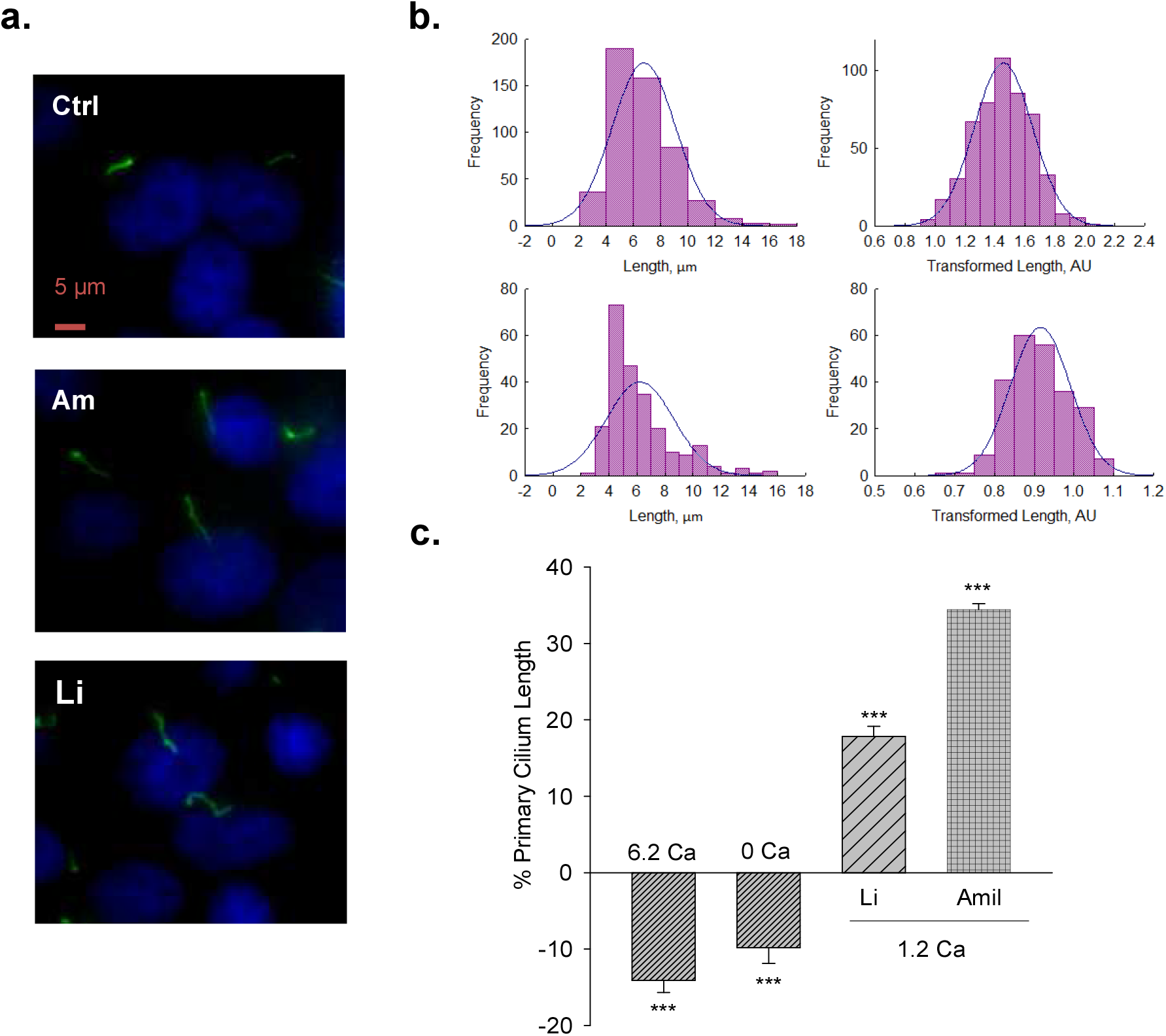
Effect of PC2 channel inhibitors on the length of the primary cilium of wild type LLC-PK1 cells. **a.** Acetylated-α-tubulin antibody immunolabeling of LLC-PK1 cells (FITC, Green) to observe the length of primary cilia under control (Ctrl, Top Panel) condition, and the presence of either amiloride (200 μM, Middle Panel), or LiCl (10 mM, Bottom Panel). Cells were incubated in normal 1.2 mM Ca^2+^. Images were obtained with x60 objective. **b.** Histograms of ciliary length measurements in normal Ca^2+^ and the presence of either amiloride (Top) or LiCl (Bottom) are shown before (Left), and after (Right), Box-Cox transformation. A not Normal left-skewed distribution of data is observed before transformation. **c.** Bar graphs of percentage increase (positive bars) or decrease (negative bars) in the length of the primary cilium under various conditions, including Free- and high (6.2 mM) Ca^2+^ external condition, as well as incubations with LiCl (10 mM) and amiloride (200 μM) respect to 1.2 Ca condition. Li^+^ and amiloride were added in the presence of external 1.2 mM Ca^2+^. All data are the mean ± SEM of percentage change as compared with respect to the control condition in normal Ca^2+^. Asterisks indicate statistical significance: *** p < 0.001 vs. control, as evaluated by one-way ANOVA.

### Effect of Li^+^ on primary cilia length

To further explore the effect of PC2 inhibition on the primary cilia length, cells were also exposed to Li^+^ (10 mM), which increases ciliary length in neurons and other cells,^30^ and has a strong inhibitory effect on PC2 channel function.^34^ The incubation procedure was similar to that used for amiloride, where the medium in this case was instead supplemented with 10 mM LiCl (Fig. 3a). Exposure of cells to Li^+^ also induced a statistically significant 17.43% ± 1.48% increase in primary cilia length (5.55 ± 0.07 μm, n = 240 vs. 4.72 ± 0.05 μm, n = 510, p <0.001, Fig. 3b & c).

### Effect of inhibition of PC2 expression on primary cilia length

To further confirm the effect of PC2 function on primary cilia length, its expression was reduced by specific PC2 siRNA transfection, as recently reported.^27^ Transfection efficacy was evaluated using a fluorescent silencing probe. PC2-silenced cells (P1ss) in normal Ca^2+^ (Fig. 4a) had statistically significant 10.25 ± 0.88% longer primary cilia as compared to cells transfected with scrambled RNA (Irss) (4.66 ± 0.04 μm, n = 1116 vs. 4.23 ± 0.04 μm, n = 957, p < 0.001, Fig. 4b & 5). Interestingly, in the presence of high external Ca^2+^, P1ss cells showed a 24.67 ± 1.65% increase in primary cilia length as compared to their respective controls, the Irss-treated cells exposed to the same external Ca^2+^ concentration (4.14 ± 0.05 μm, n = 924 vs. 3.35 ± 0.04 μm, n = 813, p < 0.001, Fig. 4b & 5).

**Fig. 4:**
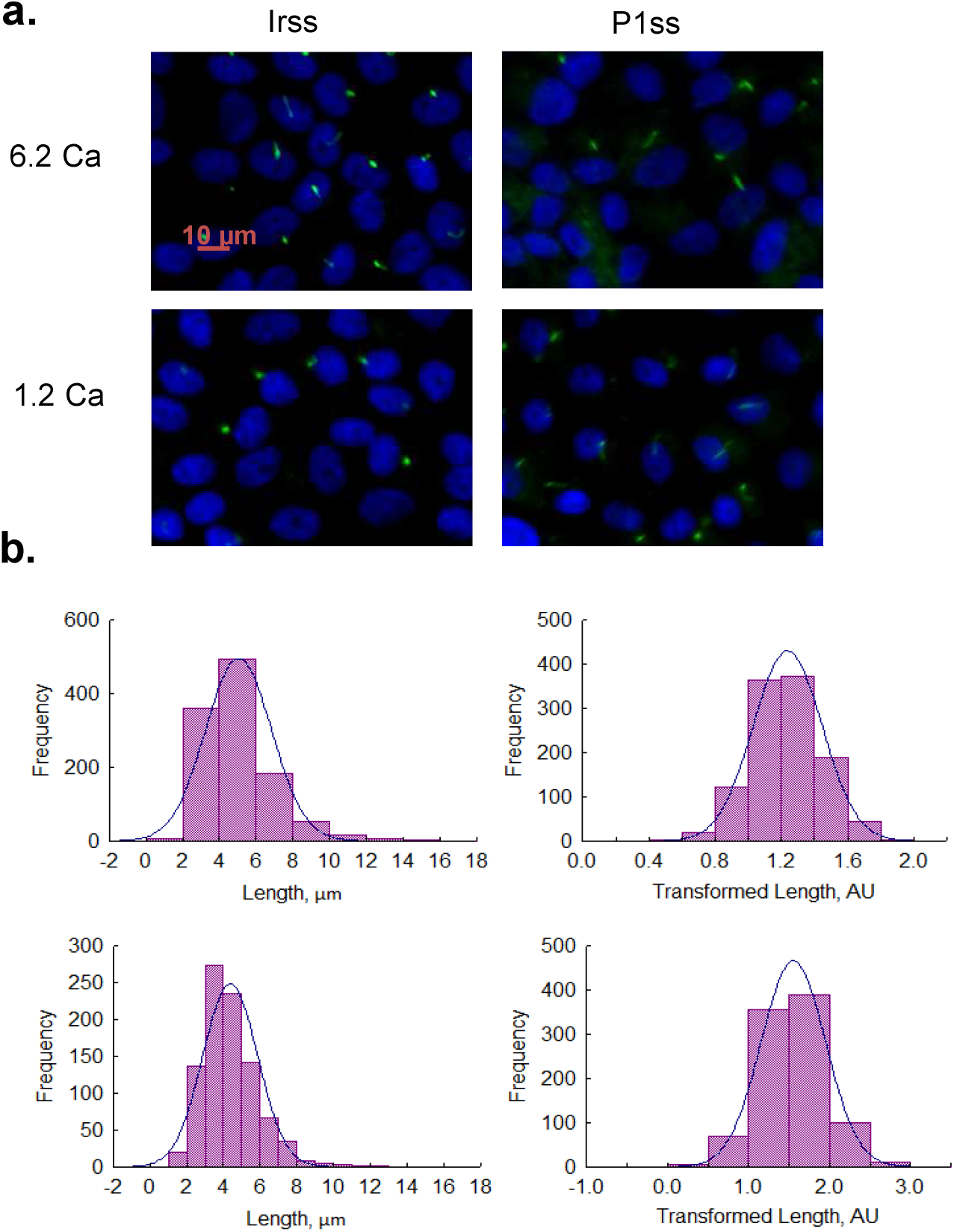
Effect of PKD2 gene silencing on the length of the primary cilium of wild type LLC-PK1 cells. **a.** Primary cilia are observed in green (FITC) and nuclei stained in blue (DAPI) in cells transfected with scrambled (Irss) and PKD2-specific (P1ss) probes in either normal (1.2 mM) or high Ca^2+^ (6.2 mM) conditions. In normal Ca^2+^, P1ss silenced cells had longer primary cilia than their respective controls (Irss, 1.2 Ca). Longer primary cilia were also observed in silenced P1ss cells in high Ca^2+^ respect to Irss, 6.2 Ca condition. **b.** Box-Cox transformation for data of the length of the primary cilium in LLC-PK1 cells after inhibition of *PKD2* gene expression under 1.2 mM (Top) and 6.2 mM (Bottom) external Ca^2+^ conditions. Frequency data distributions are shown before (Left) and after (Right) Box-Cox transformation.

**Fig. 5:**
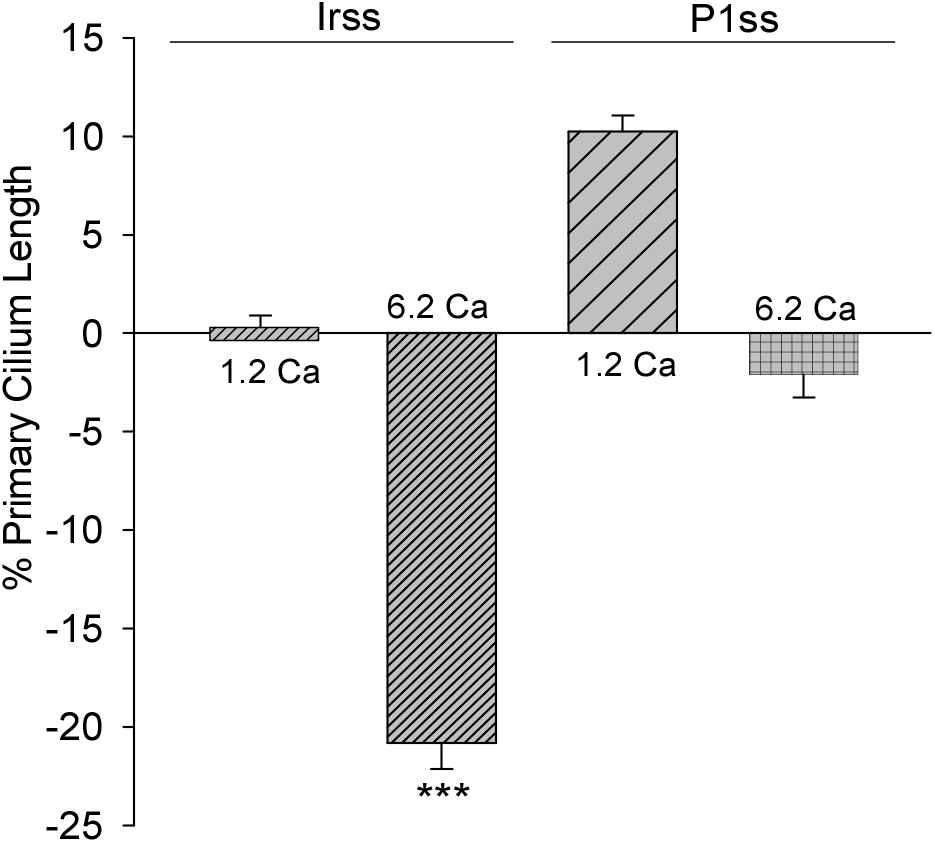
Differences in the length of the primary cilium after inhibition of PKD2 gene expression. Bar graphs represent either the percent increase (positive bars) or decrease (negative bars) of the length of the primary cilium of LLC-PK1 cells in normal (1.2 mM) and high (6.2 mM) external Ca^2+^ concentrations. Data were made relative to the Irss 1.2 mM Ca^2+^, control condition with the scrambled silencing RNA probe (Irss). Irss-treated cells responded to high external Ca^2+^ with a large decrease in the length of the primary cilium. Ciliary length of *PKD2*-silenced cells increased with respect to the it respective controls in the presence of both normal (1.2 mM) and high (6.2 mM) external Ca^2+^. Asterisks (***) indicate statistical significance at p < 0.001 vs. Irss control, evaluated by one-way ANOVA.

Altogether, the results suggest that inhibition of PC2 expression produces a primary cilium elongation in normal Ca^2+^. This phenomenon is consistent with both of the PC2 channel blockers. However, the effect of PC2 expression was Ca^2+^ dependent. In high Ca^2+^ the PKD2 silenced cells actually reversed the observed decreased in primary cilia length. Primary cilia length correlated with both the presence of a functional PC2 and the external Ca^2+^ concentration.

## Discussion

Ciliopathies are caused by defects in the formation and/or function of cilia, especially primary cilia,^35^ which render a group of clinical syndromes sharing common phenotypic features. There are more than a hundred known ciliopathies to date. Established ciliopathies include syndromes such as Bardet-Biedl (BBS), Joubert (JBTS), Meckel-Gruber (MKS), Alström, Senior-Löken, and Oro-facial-digital type 1 (OFD1), and diseases such as ADPKD and nephronophthisis (NPHP),^35^ among others, with a collective frequency of 1:1000.^36^ Since changes in primary ciliary length are considered an initial step in the formation of cysts of certain diseases, including ADPKD, in this study we explored different conditions that either control or that modulate its PC2 channel function or expression, on the length of primary cilia in LLC-PK1 cells.

There is little doubt that cilia structure, function and stability play an important role (s) in all normal kidney development and maintenance. Currently, there is a unifying hypothesis suggesting that the mechanisms of cyst formation in different genetic diseases is the result of cilia dysfunction, including abnormalities in cilia structure, composition, and signaling.^37, 38^ The vast majority of mutated proteins linked to renal cystic diseases can be localized to the primary cilium or its associated structures such as the basal body, centrosomes or ciliary transition zone.^3^ However, it remains to be determined how the “cilia hypothesis” can explain the highly variable renal phenotypic spectrum observed in different diseases.^26^ In renal tubular epithelial cells, primary cilia are thought to function as flow sensors that are required for normal epithelial proliferation and differentiation in the kidney.^39, 40^ Dysfunctional renal primary cilia change the patterns of proliferation and differentiation, resulting in cystic kidney diseases.^1, 10, 39^ The length of primary cilia increases in murine-injured kidneys,^41–43^ and in human renal transplants that suffer acute tubular necrosis.^43^ Cobalt chloride treated renal epithelial cells show much longer cilia than control cells. The hipoxia-simulating maneuver in the kidney seems to stabilize the hipoxia-inducible factor alpha (HIF-1α) transcription factor. It is speculated that elongation of primary cilia following renal injury may actually increase their sensory capacity, which is crucial to epithelial differentiation during renal repair.

The localization of the ADPKD proteins, PC1 and PC2, to primary cilia was considered an important link between primary cilia and renal cysts.^2^ It is interesting to note, however that ciliary length is normal in cells and organisms carrying *PKD1* and *PKD2* mutations, implying a defect in function rather than structure.^13^ This was supported by the observation that *PKD1* and *PKD2* mutants in the nematode *Caenorhabditis elegans* with impaired male mating behavior also have defects in cilia.^44^ The demonstration that the mammalian PC1 and PC2 proteins in renal epithelia are essential for transducing cilia-based mechanosensory Ca^2+^ signals confirmed that polycystin-mediated ciliary function was also likely to apply to epithelial cell behavior.^23^ In the particular case of ADPKD it remains an important point to explore and find functional-structural connections between PC2 and ciliary parameters. However, it is still difficult to assess ciliary function *in situ*, mainly due to the physical constraints of this organelle.

To date, methods to measure primary cilia length in cultured cells are prone to the bias of human selection, mainly caused by the orientation of the primary cilia with respect to the imaging plane.^45^ Despite the fact that confocal fluorescent imaging is the most common technique to make these measurements, primary cilia are randomly oriented within the microscopic plane. In cultured cells, for example, most primary cilia protrude more or less perpendicularly to the cell surface, and may or may not be visible in the focal plane, thus leading to inaccurate measurements.^46^ Thus, most studies focus on measuring either flat primary cilia or their projection, which risk an underestimation of the actual length. Corrections to alleviate this problem could be made based on the Pythagorean Theorem. However, this approach must assume a continuous slope along the entire length of the cilium.^45^ Although it is also possible to make a 3D reconstruction of the primary cilium to measure its length, this, instead, is a very laborious technique that requires specific software, the number of primary cilia that can be examined is usually small, and poses a significant technical challenge.^45, 46^ The present study has somewhat overcome the problem by means of increasing the number of individual observations under each experimental condition and transformation of the the data to normalize and improve the statistical validity for comparison by quantitative statistical methods. Thus, given that the length of the primary cilium is a continuous quantitative variable, and that the sample size was sufficiently large, we were able to convert the actual data to reach a Normal distribution for each group through Box-Cox transformation (see Supplemental Material).^32^ The corrected data were then compared by one-way ANOVA rendering statistically significant differences for the changes in the length of primary cilia observed under the various experimental conditions explored.

The present study explored and reconcilied data on PC2 function and expression, with the length of primary cilia in renal epithelial cells. The simplest possible working hypothesis is that being PC2, a Ca^2+^ permeable ion channel, its function might control the delivery and concentration of ciliary Ca^2+^, which in turn is a potent microtubular (MT) depolymerizing agent.^47^ Thus, we considered the possibility that PC2-mediated ciliary Ca^2+^ transport would actually contribute to modify the length of primary cilia. High external Ca^2+^ in incubation of LLC-PK1 cells resulted in the shortening of primary cilia, which is consistent with the above hypothesis that PC2-mediated an increase in intraciliary Ca^2+^ enabled the catastrophic depolymerization of axonemal MTs. It is important to note, however, that cells incubated in the absence of external Ca^2+^, also displayed shorter primary cilia (data not shown). While there is no apparent explanation for this phenomenon, it is possible that cells exposed to a Ca^2+^-free environment for several hours would also show other changes, including the loss of adhesiveness to the substrate, which could lead to the retraction of the primary cilia.

The depolymerizing effect of Ca^2+^ on MTs^47, 48^ would control the equilibrium between the assembly and disassembly of the axoneme thus regulating ciliary and flagellar length. Therefore, ciliary length was tested by PC2 inhibition with amiloride, a diuretic that increases renal excretion of Na^+^ while decreasing K^+^ excretion.^49^ Amiloride and its derivative Benzamil are high-affinity blockers of the epithelial Na^+^ channel, ENaC, acting as a blocking agent of the channel’s pore.^49^ Amiloride also functions as an inhibitor of other Na^+^ transporters and non-selective cation channels, including PC2.^15^ LLC-PK1 cells were incubated with 200 μM amiloride. The results of the treatment with amiloride at a concentration that is expected to produce full inhibition of the PC2 channel,^15^ showed an increase in ciliary length. It is also possible however, for amiloride to also block other Ca^2+^ entry mechanisms, and/or possibly, other channel such as ciliary ENaC^24^ that also affect ciliary length independently. However, LLC-PK1 cells incubated with LiCl showed primary cilia statistically longer with respect to the untreated cells, in agreement with other studies where Li^+^ treatment induced an increased in ciliary length various cell models.^30^

Treatment with Li^+^ is an important pharmacotherapy as a proven agent in bipolar disorder and mania and more recently, in psychoses and in various neurodegenerative disorders.^50^ Although, it is not yet clear as to how Li^+^ ion exerts its biological effect(s), it has been observed a lengthening of the primary cilium in various cultured cells, including neurons and fibroblasts.^30^ The primary cilium is a preferential location for PC2 cell expression,^23, 24^ and we have shown that Li^+^ reduces PC2 channel conductance and modifies the reversal potential of the *in vitro* translated PC2.^34^ Even low concentrations of Li^+^ (1-10 mM) have a considerable effect on PC2 function,^34^ suggesting the potential relevance of this ion in a clinical setting. At least two mechanistic aspects should be considered in this regard. Li^+^ could be transported to the ciliary compartment, even at low concentrations, accumulating and exerting a still unknown structural effect on the ciliary length. Another possibility is to consider the potential effect of the Li^+^ interaction with PC2 that would affect Ca^2+^ transport. In any case, Li^+^ blockade of PC2 in organelles such as the primary cilium, may help explain the impaired sensory function and the therapeutic effects of Li^+^.

To confirm the functional role of PC2 in the regulation of the length of the primary cilium, *PKD2* gene expression was silenced in the LLC-PK1 cells. In the presence of normal Ca^2+^, *PKD2*-silenced cells showed a longer primary cilium as compared to their respective controls. This is consistent with the fact that the absence of PC2 would have a similar effect to that of PC2 inhibitors, giving rise to longer primary cilia and suggesting that PC2 is necessary for the regulatory Ca^2+^ entry into the organelle. Interestingly, in the presence of high external Ca^2+^, PC2-silencing induced a reversal such that the values were similar to the control cells incubated in normal Ca^2+^. Although there is no explanation for this phenomenon, it may be related to the fact that PC2 silencing only represented approximately 50% of the inhibited gene product, such that it is possible that the remaining gene product could compensate and maintain a sustainable Ca^2+^ influx to the primary cilium. In the absence of PC2 Ca^2+^ transport could also be potentiated by other Ca^2+^-permeable channels present in the primary cilium, including TRPC1 and TRPV4.^24, 51^ In fact, PC2 interacts with these TRP channel isotypes form different hetero-multimeric complexes. PC2/TRPC1 hetero-multimeric channels, for example, show different conductance patterns than their homomeric counterparts,^52^ and may modify their Ca^2+^ permeability properties, although information in this regard is yet unavailable. PC2 silencing, could possibly affect these interactions, thus interfering with the mechano-transduction of the Ca^2+^ signals associated with primary cilia.

In conclusion, our results indicate that the length of the primary cilium in renal epithelial cells is controlled by a functional PC2, such that pathways that lead to its inhibition, including gene suppression, and possibly a consequent decrease in ciliary Ca^2+^ entry, render ciliary elongation, while the presence of high external Ca^2+^ leads to a decrease in its length (Fig. 6). These results open the possibility to elucidate the relationship between changes in ciliary length and Ca^2+^ influx in the onset of ciliopathies and the formation of signals leading to the formation of renal cysts, more specifically, in ADPKD.

**Fig. 6:**
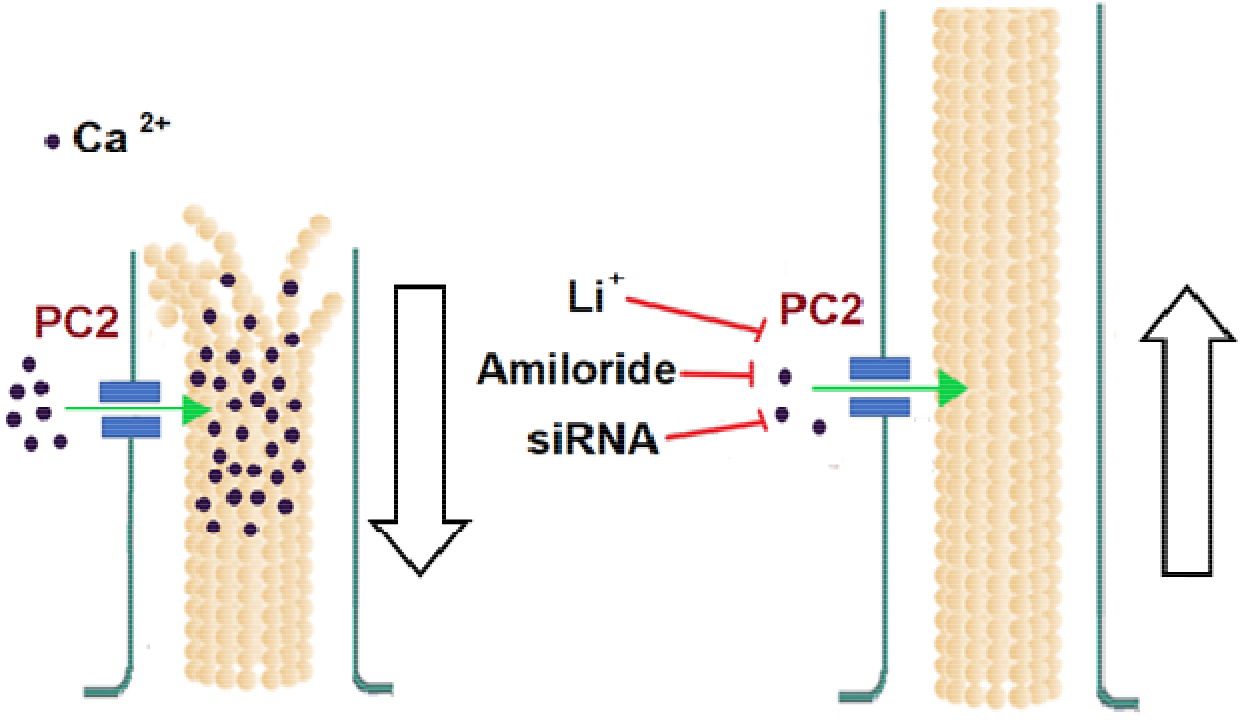
Proposed model of primary cilium length regulation by PC2. Cartoon of the possible role(s) PC2 plays in the regulation of the length of the primary cilium in LLC-PK1 renal epithelial cells. Under physiological external Ca^2+^ conditions (1.2 mM), the length primary cilium length is maintained by PC2-mediated Ca^2+^ entry. In the presence of a high external Ca^2+^ (6.2 mM) concentration, the increased intraciliary ion would help depolymerize axonemal microtubules, leading to reduction in their length, and thus that of the primary cilium. Either PC2 inhibition (by Li^+^ or amiloride) or reduction of its expression (siRNA), would instead contribute to reducing Ca^2+^ entry to the primary cilium, also reducing the rate MT depolymerization, rendering longer primary cilia.

## Author contributions

P.L.P. and N.S. carried out all experimental procedures, conducted the analysis of the experimental data, and prepared Figures. H.F.C. and M.R.C. designed all the experiments and wrote the main manuscript text. All authors approved the final version of the manuscript.

## Acknowledgments

The present study was funded by grant PICT 2012 N°1559 (HFC), MinCyT-Argentina. The authors do not have financial interests.

## Supplemental Material

The statistical analyses of the experimental data were as follows. Data from -either three or- four experiments were collected for any given condition (in the present example two conditions), 1.2 mM and 6.2 mM external Ca^2+^ concentrations. Data were shown to be wide variable for the ciliary length measurements, even when cells were grown and kept undertissue culture conditions until total confluence. This variability made difficult any comparisons among groups. The incubation in high Ca^2+^ produced a decrease in ciliary length as compared to cells kept in normal Ca^2+^ (1.2 mM). This trend was observed for all four repeats (Exps 1-4, See Table A1, and Fig. 1a). Because no data followed a Normal distribution, the non-parametric one-way ANOVA test was first performed by Kruskall-Wallis ranges, for the 1.2 and 6.2 mM Ca^2+^ conditions. The results indicated that in normal Ca^2+^, experiments 3 and 4 were not significantly different, but the others were (p < 0.001). In high Ca^2+^, in contrast, no significant differences were found between experiments 1 and 2 and between experiments 3 and 4. Table A1 shows the median and the 25% and 75% quartiles obtained under both conditions for each one of the experiments.

**Table A1.**
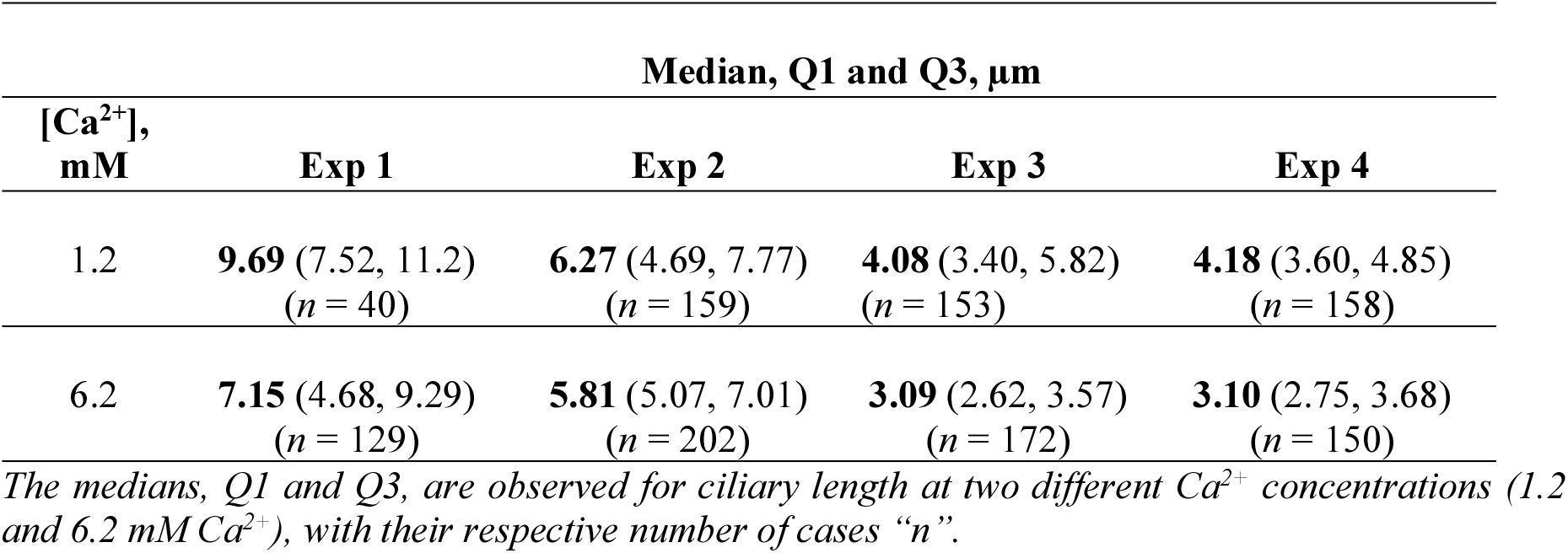
Ciliary length values for cells exposed to different concentrations of external Ca^2+^. Data are the medians and the 25% and 75% quartiles.

| [$\text{Ca}^{2+}$],<br>mM | Median, Q1 and Q3, $\mu\text{m}$ | | | |
| --- | --- | --- | --- | --- |
|  | Exp 1 | Exp 2 | Exp 3 | Exp 4 |
| 1.2 | **9.69** (7.52, 11.2)<br>($n = 40$) | **6.27** (4.69, 7.77)<br>($n = 159$) | **4.08** (3.40, 5.82)<br>($n = 153$) | **4.18** (3.60, 4.85)<br>($n = 158$) |
| 6.2 | **7.15** (4.68, 9.29)<br>($n = 129$) | **5.81** (5.07, 7.01)<br>($n = 202$) | **3.09** (2.62, 3.57)<br>($n = 172$) | **3.10** (2.75, 3.68)<br>($n = 150$) |

*The medians, Q1 and Q3, are observed for ciliary length at two different $\text{Ca}^{2+}$ concentrations (1.2 and 6.2 mM $\text{Ca}^{2+}$ ), with their respective number of cases “n”.*

To assess statistical differences between experimental conditions (in this example the two Ca^2+^ concentrations) for each particular experiment, the Mann-Whitney U test was used instead. The results showed significant differences between experiments 1, 3 and 4 (p < 0.001), but not for experiment 2 (data not shown). Thus, we sought to normalize the data groups under each experimental condition, in order to apply parametric tests.

To be able to compare means and Standard Errors between groups and the subsequent statistical analysis with more powerful tests, data were normalized by Box-Cox transformation (see main text, Figs. 1–4). From the transformation formula (see Materials & Methods), the value of the mean with its SEM was obtained for each incubation condition. The one-way ANOVA parametric test was then performed. Table A2 shows the mean ± SEM for normalized values under both Ca^2+^ concentrations. For the 1.2 Ca^2+^ condition, only experiments 3 and 4 showed no significant differences between each other, as with the Kruskall-Wallis test. In the 6.2 mM Ca^2+^ condition, experiments 3 and 4 showed no significant differences, but differences were observed for all others (p < 0.001).

**Table A2.** Ciliary length values for cells exposed to different concentrations of external Ca^2+^. Data are the mean ± SEM after Box-Cox re-transformation of original data.

| [$\text{Ca}^{2+}$],<br>mM | Mean ± SEM | | | |
| --- | --- | --- | --- | --- |
|  | Exp 1 | Exp 2 | Exp 3 | Exp 4 |
| 1.2 | 9.66 ± 0.22<br>(n = 40) | 6.12 ± 0.11<br>(n = 159) | 4.08 ± 0.10<br>(n = 153) | 4.18 ± 0.10<br>(n = 158) |
| 6.2 | 6.96 ± 0.25<br>(n = 129) | 5.83 ± 0.08<br>(n = 202) | 3.03 ± 0.08<br>(n = 172) | 3.12 ± 0.09<br>(n = 150) |

*Means ± SE were observed at different concentrations of $\text{Ca}^{2+}$ (1.2 and 6.2 mM $\text{Ca}^{2+}$ ) with their respective n.*

Once the data were re-transformed, the tendency to reduce ciliary length after incubation in 6.2 Ca again was observed. This was evident in the four experiments (Fig A1). The Student *t* test statistic showed significant differences for each of the comparisons between the 1.2 Ca and 6.2 Ca conditions (p < 0.03 for experiment 2 and p < 0.001 for experiments 1, 3 and 4).

**Fig A1.**
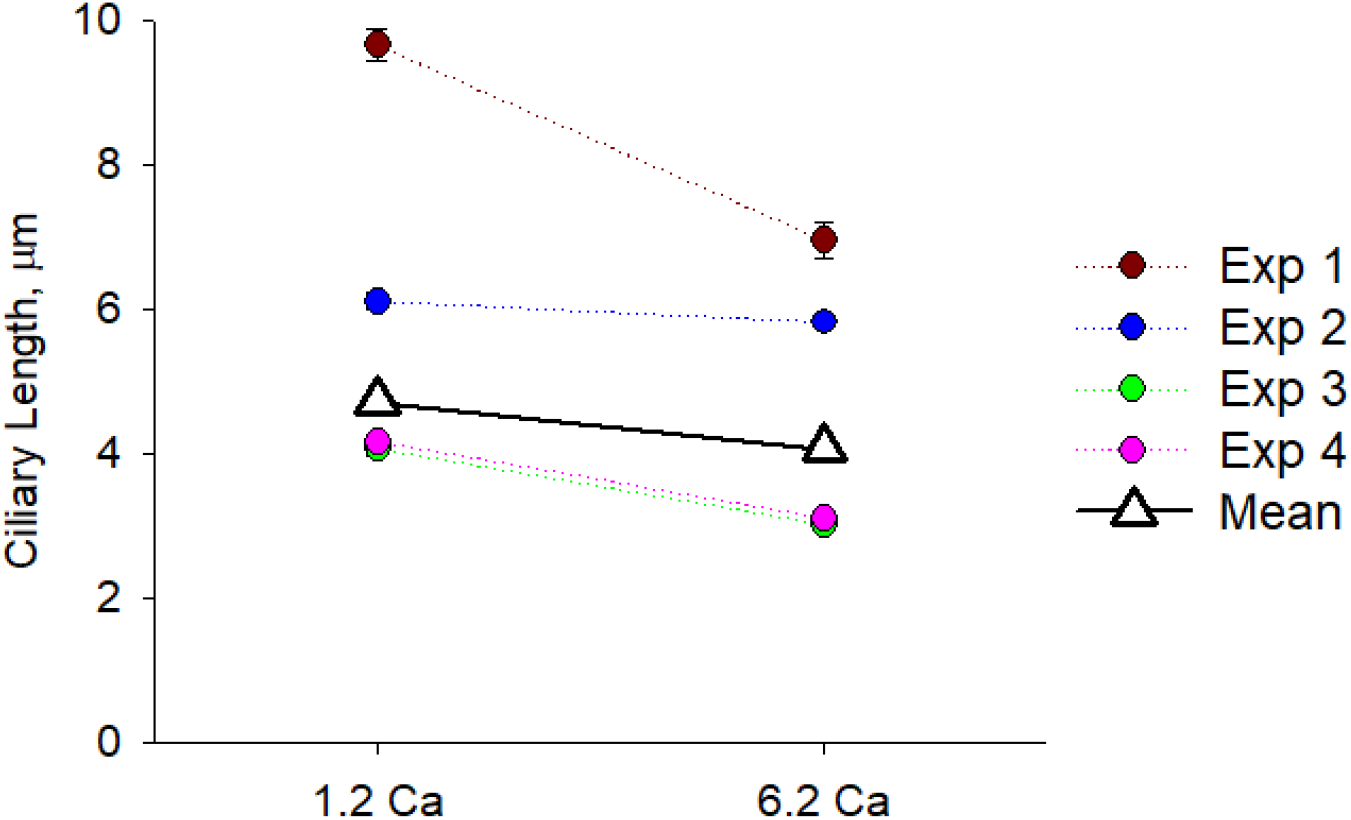
Effect of external Ca^2+^ on ciliary length for each one of the four experiments after Box-Cox re-transformation of original data. Statistical difference of individual experiments evaluated by Student t test was as follows, exps, 1, 3, and 4 p < 0.001, and exp 2, p < 0.05, with respect to the control condition. Please note that open triangles represent the average for the four experiments for both conditions. Statistical significance is indicated in the text.

Fig. A1 shows (open triangles) pooled average of normalized data for the four experiments under each condition, rendering the values reported in the manuscript, 4.72 ± 0.05 μm, n = 510 vs. 4.08 ± 0.06 μm, n = 653, p < 0.001, for the 1.2 and 6.2 mM Ca^2+^, respectively. In the same way that incubating the cells in 6.2 mM Ca^2+^ produced a reduction of primary cilia length, which is reported by the pooled data, a similar approach was conducted for allremaining conditions. Namely, after evaluating each of the experiments among themselves, values were included for each condition in a common pool after Box-Cox transformation, using mean ± SEM for the subsequent application of parametric statistical tests. All experimental groups showed strong Normalization and a significant statistical difference as compared to its own respective control condition (see main manuscript).

